# A biofilm activator MmtA1 modulates multi-drug resistance in mycobacteria

**DOI:** 10.1101/2025.04.03.647061

**Authors:** Jialing Hu, Chun-Hui Gao, Min Yang, Wei Wang

## Abstract

When exposed to the antibiotic-induced stress, *Mycobacterium* species have the ability to survive by forming protective multicellular structures known as biofilms. However, the underlying mechanism of biofilm formation in response to antibiotics is not fully understood. In this study, we identified a transcriptional factor, MmtA1, which responds to the first-line anti-tuberculosis drug isoniazid and regulates genes involved in sugar and lipid transportation. This leads to increased biofilm formation and multi-drug resistance in *M. smegmatis*. These findings suggest that isoniazid may act as a molecular messenger, working in coordination with transcriptional regulatory factors to induce multidrug resistance under chemotherapy.

**Importance:** This study elucidates a novel transcriptional regulatory mechanism by which *Mycobacterium smegmatis* develops multidrug resistance in response to the anti-tuberculosis drug isoniazid. By linking antibiotic stress to transcriptional regulation and metabolic adaptation, this work provides new insights into drug-induced tolerance in mycobacteria, suggesting antibiotic functional as inducer of biofilm formation, triggering a coordinated transcriptional response to enhance bacterial survival under antibiotic pressure.

## Introduction

Biofilm formation is the process by which a group of planktonic, free-living bacterial cells convert into a complex, surface-attached microorganism community, providing a physical barrier to environmental stresses (1, 2). Thus, the ability to form biofilms affects bacterial persistence in response to stress conditions (1, 3, 4). Most species of mycobacteria have a strong tendency to form biofilm in liquid culture (5). For pathogenic *Mycobacterium tuberculosis*, pellicle biofilms have been shown to increase its drug tolerance *in vitro* (6). During infection, biofilm formation in lungs has been shown to protect bacilli from the host immune system and antimycobacterial agents, allowing for the emergence of a drug-tolerant phenotype as well (7). Recent evidence suggests that, the bactericidal rate of either isoniazid or rifampicin against *M. tuberculosis* in the biofilm state is reduced compared to the bacteria in the planktonic form (6). Environmental mycobacteria *Mycobacterium smegmatis* biofilms are also reported to harbor more drug-tolerant persisters than planktonic cultures (8, 9). Therefore, it is critical to recognize the importance of the general characteristics and putative regulatory mechanism of mycobacterial biofilms formation.

Antibiotics, particularly at low doses, can serve as biofilm modulators in bacteria. When exposed to exogenous antibiotics, the stimulation of bacterial biofilm formation is frequently observed (10). For example, *Pseudomonas aeruginosa* was reported to increase biofilm production when challenged with sub-inhibitory tobramycin concentrations (11). Recent advances in concept of antibiotics as signaling molecules have provided new insights into their role in biofilm regulation. Mechanistically, this phenomenon involves coordinated regulation between stress responses and extracellular matrix (ECM) synthesis in bacteria, supporting antibiotic stressed environment leading to the bacterial biofilm formation (12). The evolutionary advantages of the biofilm-mediated protection enhanced antibiotic resistance in bacteria while simultaneously creating a structural ECM barrier in biofilm and provide protection against other environmental threats (13). However, the mechanism underlying antibiotic-induced biofilm formation remains poorly characterized in mycobacteria.

Mycobacteria within biofilm communities are associated with a complex architecture of extracellular polymetric substances containing imported compounds, sugars, lipids and amino acids (14–16). In our previous study, we found that the transcriptional factors MmtA1 (Ms0179), MmtA2 (Ms0180) and Ms5575 cooperatively regulated mycobacterial biofilm formation by increasing sugar accumulation to bacterial surface in *M. smegmatis* (17). A mannitol metabolism and transportation gene cluster, *mmt* operon, was found to be the target of MmtA2, and played an essential role in the biosynthesis and secretion of D-mannitol.

In this study, we discovered that the transcription factor MmtA1 is induced by isoniazid and is associated with antibiotic resistance in M. smegmatis due to its regulatory function. Moreover, we investigated the two adjacent transcriptional regulators, MmtA1 and MmtA2, and found that they promote mycobacterial biofilm formation by positively regulating the *mmt* operon via specific DNA motifs. By overexpressing MmtA1, we identified several differentially expressed genes, including those encoding for sugar and lipid transporters, which may be responsible for mycobacterial biofilm formation. Our results demonstrate that isoniazid acts as a molecular messenger and coordinates with the transcriptional regulatory pathway in response to antibiotic pressure, leading to drug-induced tolerance of mycobacteria under antibiotic stress.

## Results

### Transcriptional regulator MmtA1 contributes to mycobacterial INH resistance

In a previous study, we characterized a regulatory pathway for biofilm formation in *M. smegmatis*. The pathway is coordinated by the *mmt* operon and its activators MmtA1 (Ms0179) and MmtA2 (Ms0180) (Fig. 1A). Further results showed that overexpression of *mmtA1* resulted in a shifted minimum inhibitory concentration (MIC) in *M. smegmatis*. The MIC of the *mmtA1*-overexpressing strain was 5.73 μg/cm^2^, while the MIC of the vector control Msm/pMV261 was 4.46 μg/cm^2^ against isoniazid (Fig. 1B). This suggests that MmtA1 is responsible for regulating INH resistance in *M. smegmatis*.

**Figure 1.**
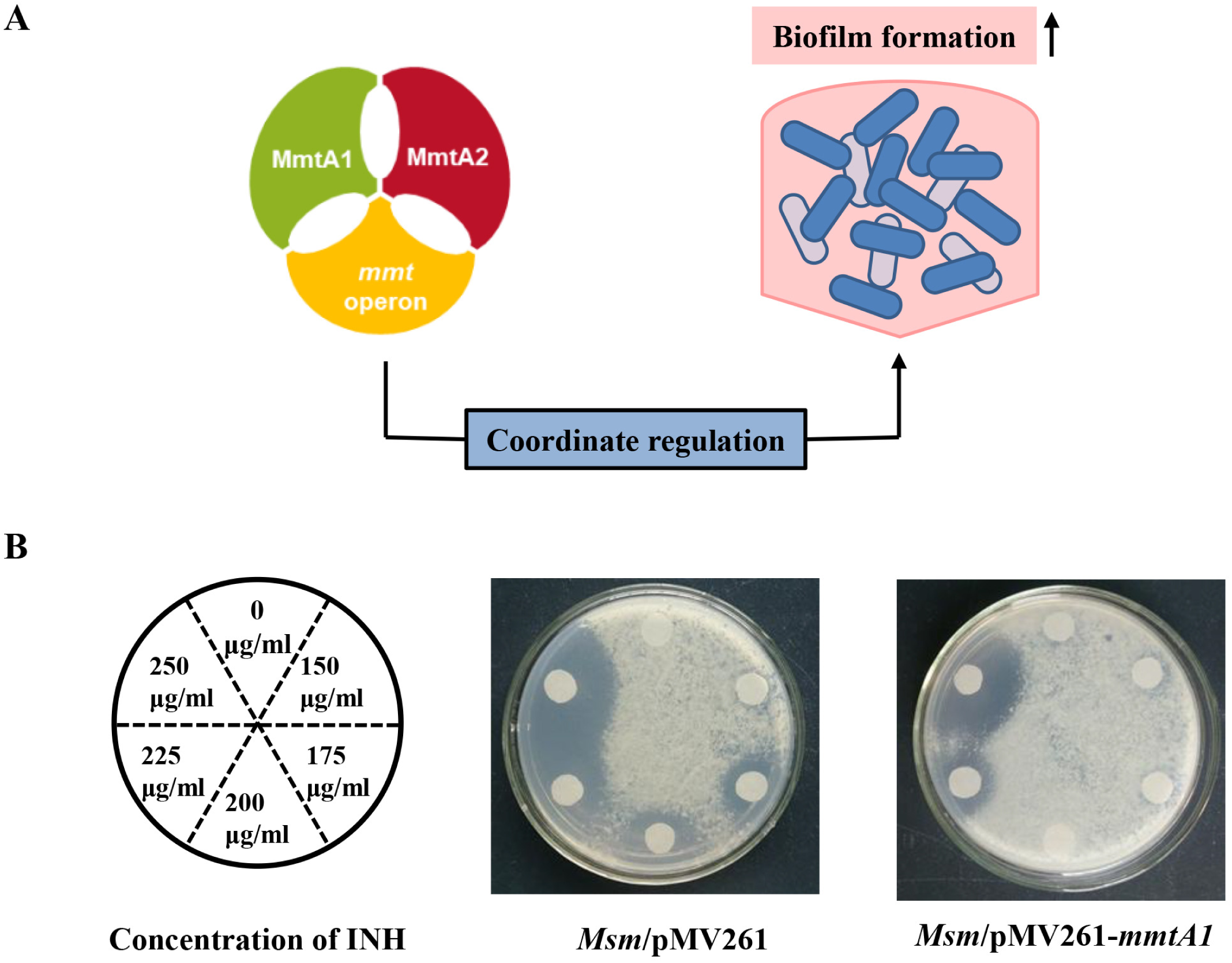
*Mycobacterium smegmatis* MmtA1 contributes to mycobacterial isoniazid resistance. (A) A general view of biofilm regulatory pathway involving the *mmt* operon and its activators in *M. smegmatis*. (B) Minimum inhibitory concentration (MIC) of isoniazid to wild-type and *mmtA1*-overexpressing *M. smegmatis* strains.

### MmtA1 and MmtA2 up-regulate multi-drug resistance in *M. smegmatis*

As described previously, MmtA1 positively regulates the expression of *mmtA2* (*Ms0180*), an adjacent gene on the *M. smegmatis* genome (Fig. 2A). To investigate the involvement of MmtA1 and MmtA2 in mediating bacterial resistance to antitubercular drugs, we examined bacterial viability under the pressure of isoniazid, rifampicin, and ethambutol. Compared with the wild-type strain, *mmtA1*- and *mmtA2*-overexpressing strains showed higher viability under INH-treated conditions but exhibited similar survival states in normal 7H9 medium without antibiotics (Fig. 2B, Fig. S1A). As shown in Fig. 2C, *mmtA1*- and *mmtA2*-deleted strains showed lower viability in the presence of INH. The expression of *mmtA1* and *mmtA2* also resulted in rifampicin and ethambutol resistance in *M. smegmatis* (Fig. 2B and 2C, Fig. S1A and S1B). These results indicate that the expression of *mmtA1* and *mmtA2* lead to multi-drug resistance in *M. smegmatis*.

**Figure 2.**
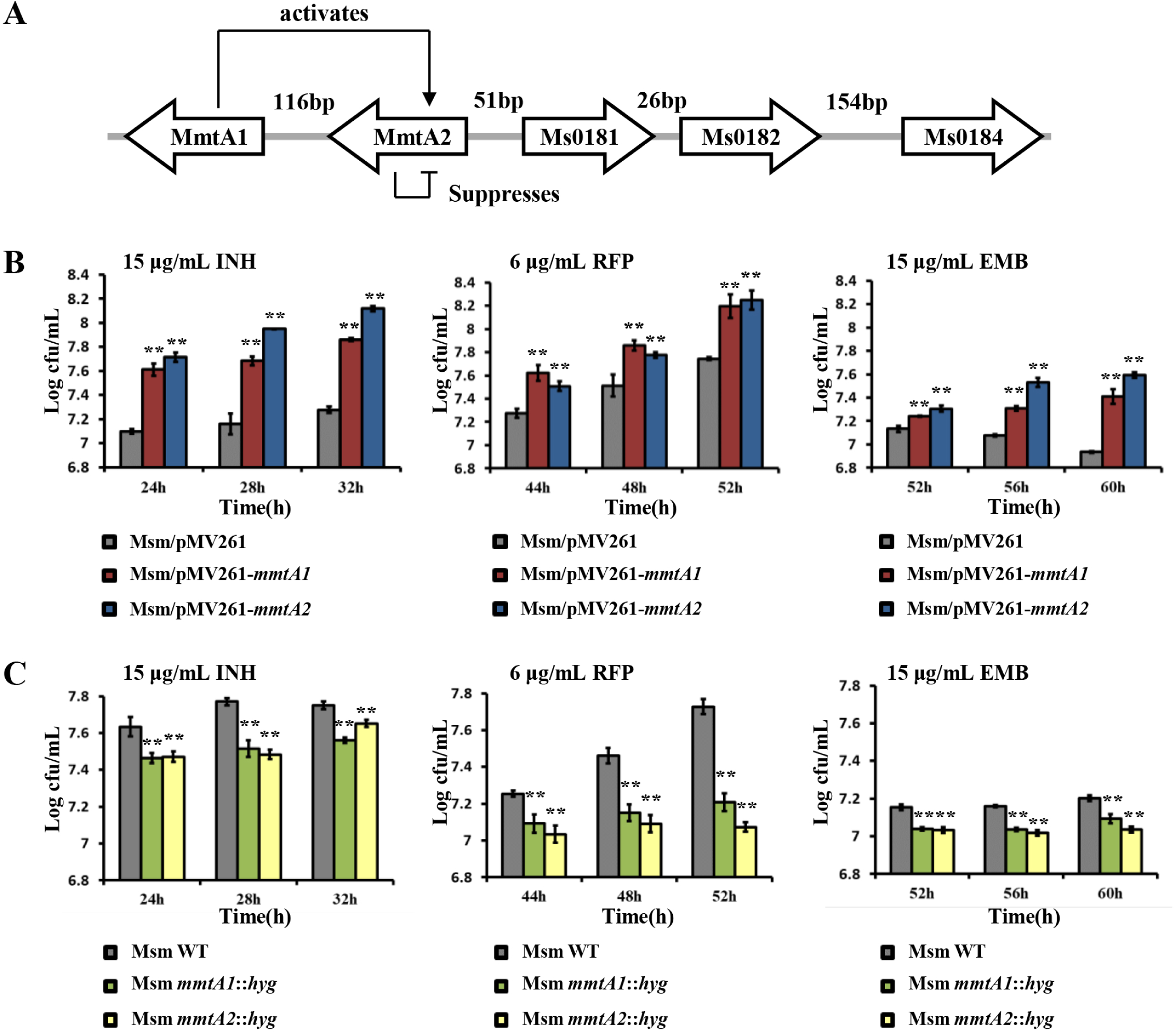
MmtA1 and MmtA2 up-regulate multi-drug resistance in *M. smegmatis*. (A) Location and transcriptional regulation of MmtA1 and *mmtA2* on *M. smegmatis* genome. (B) Drug susceptibility assays. Wild-type, *mmtA1*-overexpressing and *mmtA2*-overexpressing strains were grown in 7H9 medium supplemented with 15 μg/mL isoniazid (*left*), 6 μg/mL rifampicin (*middle*) and 15 μg/mL ethambutol (*right*). (C) Wild-type, *mmtA1-* and *mmtA2*-deleted strains were grown in 7H9 medium supplemented with 15 μg/mL isoniazid (*left*), 6 μg/mL rifampicin (*middle*) and 15 μg/mL ethambutol (*right*). *, *P* < 0.05; **, *P* < 0.01.

### MmtA1 and MmtA2 positively regulate multi-drug resistance by activating the D-mannitol related *mmt* operon

MmtA2 positively regulates the *mmt* operon, a gene cluster involved in mannitol metabolism and transportation (17). The *mmt* operon consists of genes encoding a sugar ABC transporter (*Ms5571*, *Ms5572* and *Ms5573*), a substrate-binding protein (*Ms5574*), a repressor (*Ms5575*), and a mannitol 2-dehydrogenase (*Ms5576*), which catalyzes D-mannitol metabolism (Fig. 3A).

**Figure 3.**
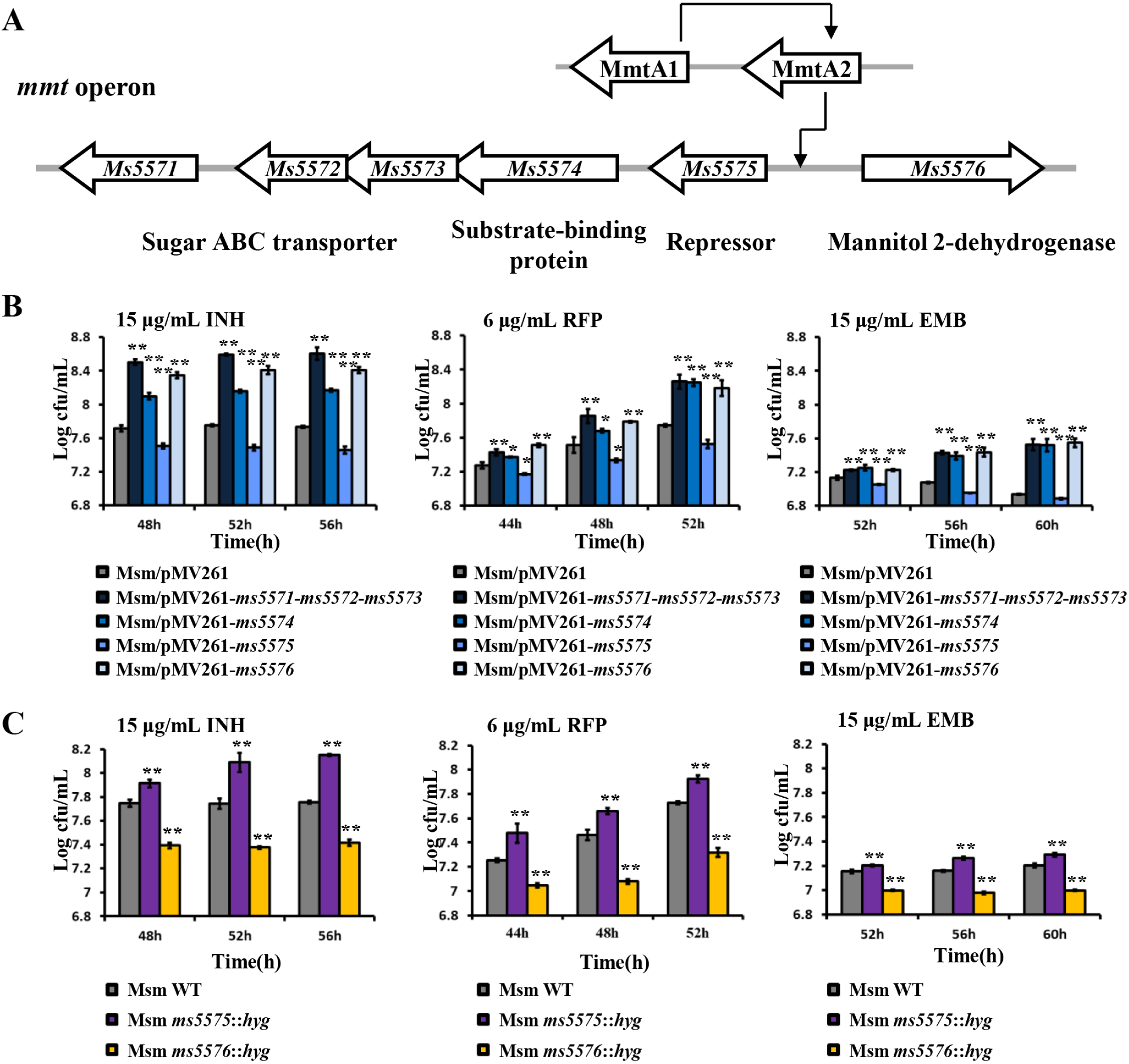
Transcriptional factors MmtA1 and MmtA2 positively regulates multi-drug resistance through the *mmt* operon. (A) The genomic location illustration of *mmt* operon. The Ms5571-Ms5575 gene cluster shares an upstream promoter region with Ms5576. (B) Drug susceptibility assays. Wild-type and *mmt*-overexpressing strains were grown in 7H9 medium supplemented of 15 μg/mL INH, 6 μg/mL rifampicin and 15 μg/mL ethambutol. (C) Wild-type and *mmt*-deleted strains were grown in 7H9 medium supplemented of 15 μg/mL INH, 6 μg/mL rifampicin and 15 μg/mL ethambutol. *, *P* < 0.05; **, *P* < 0.01.

We subsequently investigated the role of *mmt* operon in mycobacterial antibiotic resistance. When grown in a medium containing 15 μg/ml INH, most overexpressing strains exhibited higher viability compared to the wild-type (Fig. 3B). As a repressor of the *mmt* operon, Ms5575 conferred isoniazid sensitivity to *M. smegmatis* (Fig. 3B). The results indicated that the deletion of the D-mannitol biosynthesis related gene, *Ms5576*, reduced bacterial viability in the presence of INH (Fig. 3C). In contrast, deletion of the repressor Ms5575 rendered *M. smegmatis* less susceptible to isoniazid (Fig. 3C). Similar results were observed for rifampin and ethambutol (Fig. 3B and 3C; Fig.S2A and S2B). Therefore, these findings suggest that genes involved in D-mannitol metabolism and transportation enhance the antibiotic resistance of *M. smegmatis* to a variety of drugs.

### Isoniazid induces expression of MmtA1

RT-qPCR was performed to investigate the effect of isoniazid on gene expression. Compared to untreated bacteria, the expression level of *mmtA1* increased by 2.1-fold and 6.9-fold in the presence of 15 μg/ml INH and 30 μg/ml INH, respectively (Fig. 4A). However, the expression level of the negative control, *Ms0535* (18) did not exhibit any significant changes in the presence or absence of INH.

**Figure 4.**
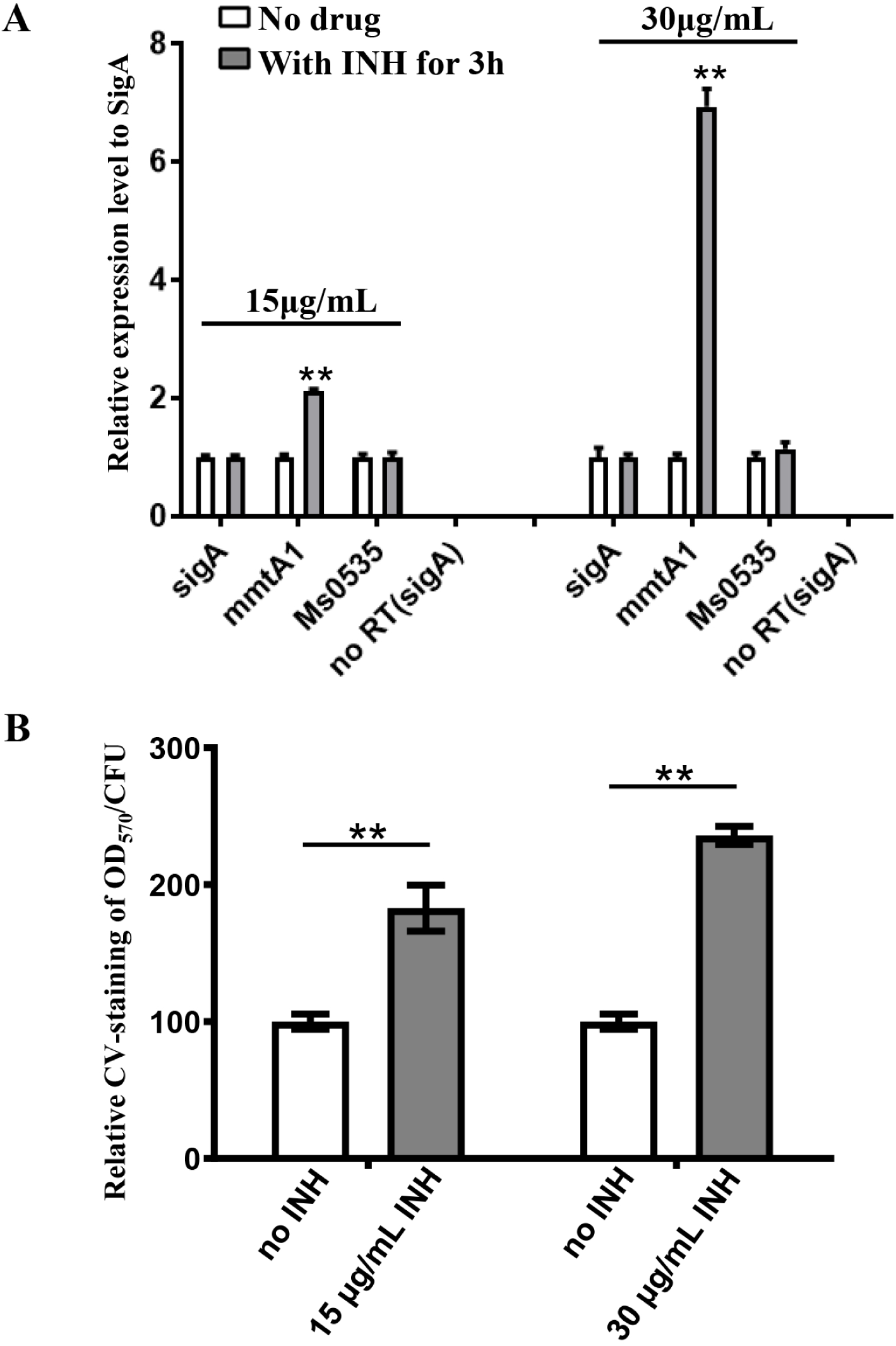
The effect of INH on the expression of MmtA1 and mycobacterial biofilm formation. (A) Quantitative real-time PCR assays for MmtA1 mRNA expressive levels under various INH concentrations. (B)Biofilm formation assays of *M. smegmatis* supplemented with INH. The columns represent the ratio of relative CV-staining and colony-forming unit (CFU) of corresponding samples. **, *P* < 0.01.

Previous studies have demonstrated that MmtA1 upregulated biofilm formation of *M. smegmatis* (17). Furthermore, our results showed that isoniazid significantly enhances biofilm formation in *M. smegmatis*. The amount of bacterial biofilm increased by 1.8-fold and 2.4-fold in the presence of 15 μg/ml INH and 30 μg/ml INH, respectively (Fig. 4B). The results suggested that isoniazid promotes *M. smegmatis* biofilm formation through the regulation manner of MmtA1.

### Transcriptional factor MmtA1 up-regulates mmtA2 by recognizing a palindromic motif

We analyzed the regulatory properties of MmtA1 by determining its binding sites to the target promoters. EMSA was performed with purified MmtA1 protein and its self-promoter MmtA1p. The result showed that MmtA1 interacts with its cognate operator DNA in a specific manner, as demonstrated by a competitive experiment performed with FITC-labeled substrate (Fig. 5A). Further analysis using the DNaseI footprinting assay identified that the binding region of MmtA1 is a two-part 16-bp palindromic DNA sequence 5’-ACAATTGT••••CCGG-3’ located from located from −68 to −53 upstream of the *mmtA1* gene (Fig. 5B). Subsequent EMSA tests suggested that the first palindrome, 5’-ACAATTGT-3’, of the identified motif is required for the MmtA1 binding (Fig. 5C).

**Figure 5.**
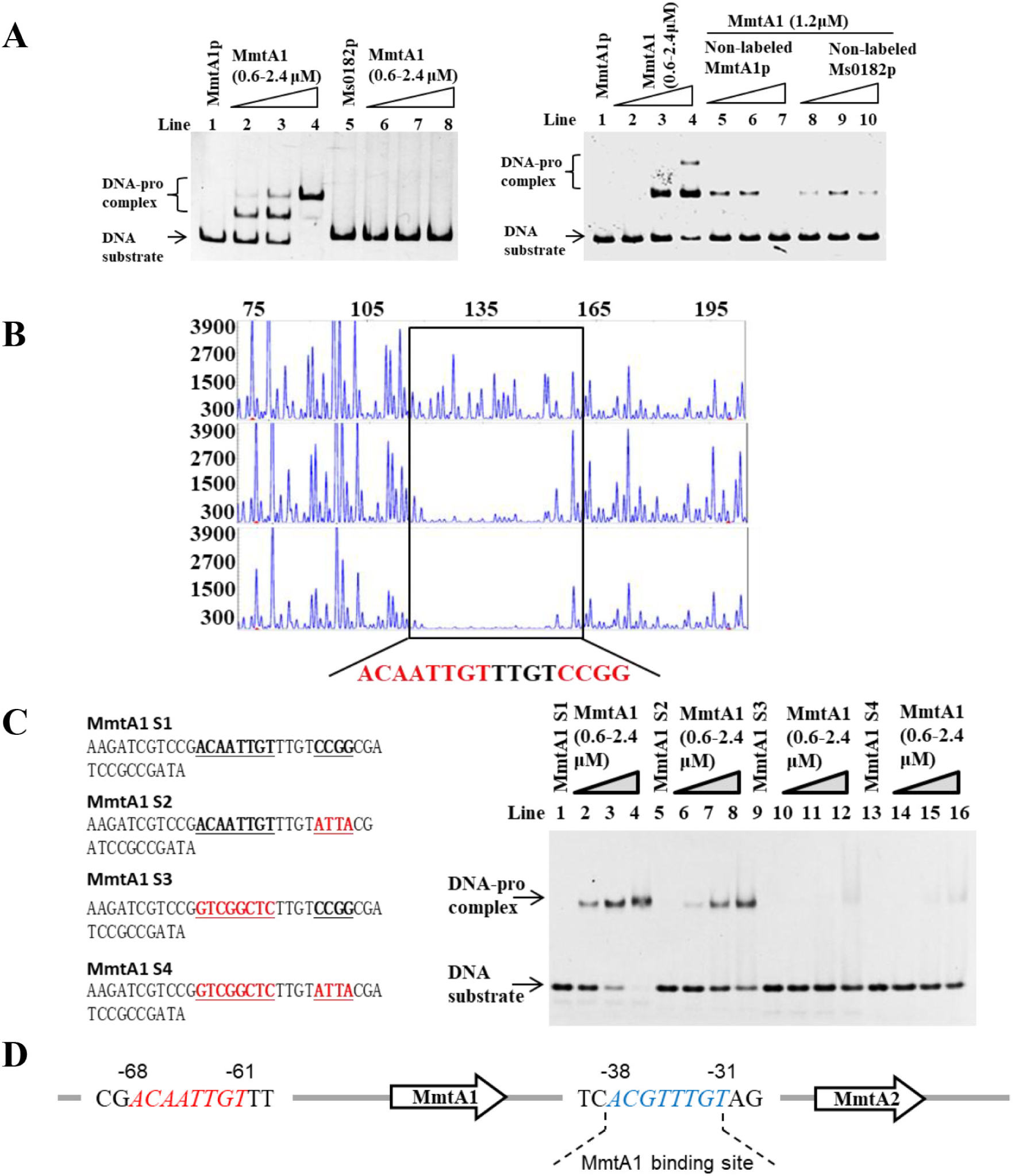
Transcriptional factor MmtA1 binds to a palindromic motif. (A) *Left*, EMSA analysis of MmtA1 binding activity to MmtA1 promoter DNA. MmtA1p (lanes 1–4) and Ms0182p (negative control, lanes 5–8) were co-incubated with increasing concentrations of purified MmtA1 protein (0, 0.6, 1.2 and 2.4 μM). *Right*, competitive binding assays for specific DNA-binding activity of MmtA1 on MmtA1p. FITC-labeled MmtA1p was pre-incubated with MmtA1 protein, followed by competition with unlabeled MmtA1p DNA (lane 5–7) and negative control Ms0182p DNA (lane 8–10). (B) DNase I footprinting assays for determination of MmtA1 binding motif. *Top*, DNA only; *middle*, 0.6 μM MmtA1; *bottom*, 2.4 μM MmtA1. (C) EMSA analysis of MmtA1 binding to DNA substrates MmtA1 S1 (lanes 1–4), MmtA1 S2 (lane 5–8), MmtA1 S3 (lane 9–12) and MmtA1 S4 (lanes 13–16). Either DNA substrate was co-incubated with 0.6–2.4 μM of MmtA1 protein. (D) Hypothetical MmtA1 binding site in the *mmtA2* promotor.

Previous experiments have demonstrated that MmtA1 activates *mmtA2* by directly binding to its promotor region (17). The potential binding motif, ACGTTTGT, which is present in the *mmtA2* promotor region (from −38 to −31 bp), suggests a possible MmtA1 binding site (Fig. 5D). β-galactosidase activity assay indicates that MmtA1 positively regulates *mmtA2* promoter via this binding motif (Fig. S3A). Taken together, the results suggested that MmtA1 up-regulates *mmtA2* expression through its binding to the palindromic motif in the *mmtA2* promoter.

### Transcriptional factor MmtA2 binds to a palindrome motif and acts as a *mmt* activator

We characterized the DNA binding activity of the HTH-like transcriptional factor MmtA2 using its self-promoter DNA, MmtA2p. EMSA revealed that MmtA2 has a specific DNA-binding activity on the DNA substrate MmtA2p (Fig. 6A). As shown in Figure 6B, there were two MmtA2 binding regions on the MmtA2p fragment. The motif sequences were confirmed as 5’-TGTAGTCGAATGACTACA-3’ and 5’-TGTAGTCGAGTGACTACA-3’ (Fig. 6B). The two binding motifs of MmtA2 share the same 14-bp palindrome regions which were recognized as 5’-TGTAGTC••••GACTACA-3’. EMSA results showed the palindrome motif is required for MmtA2 binding to DNA substrates (Fig. 6C).

**Figure 6.**
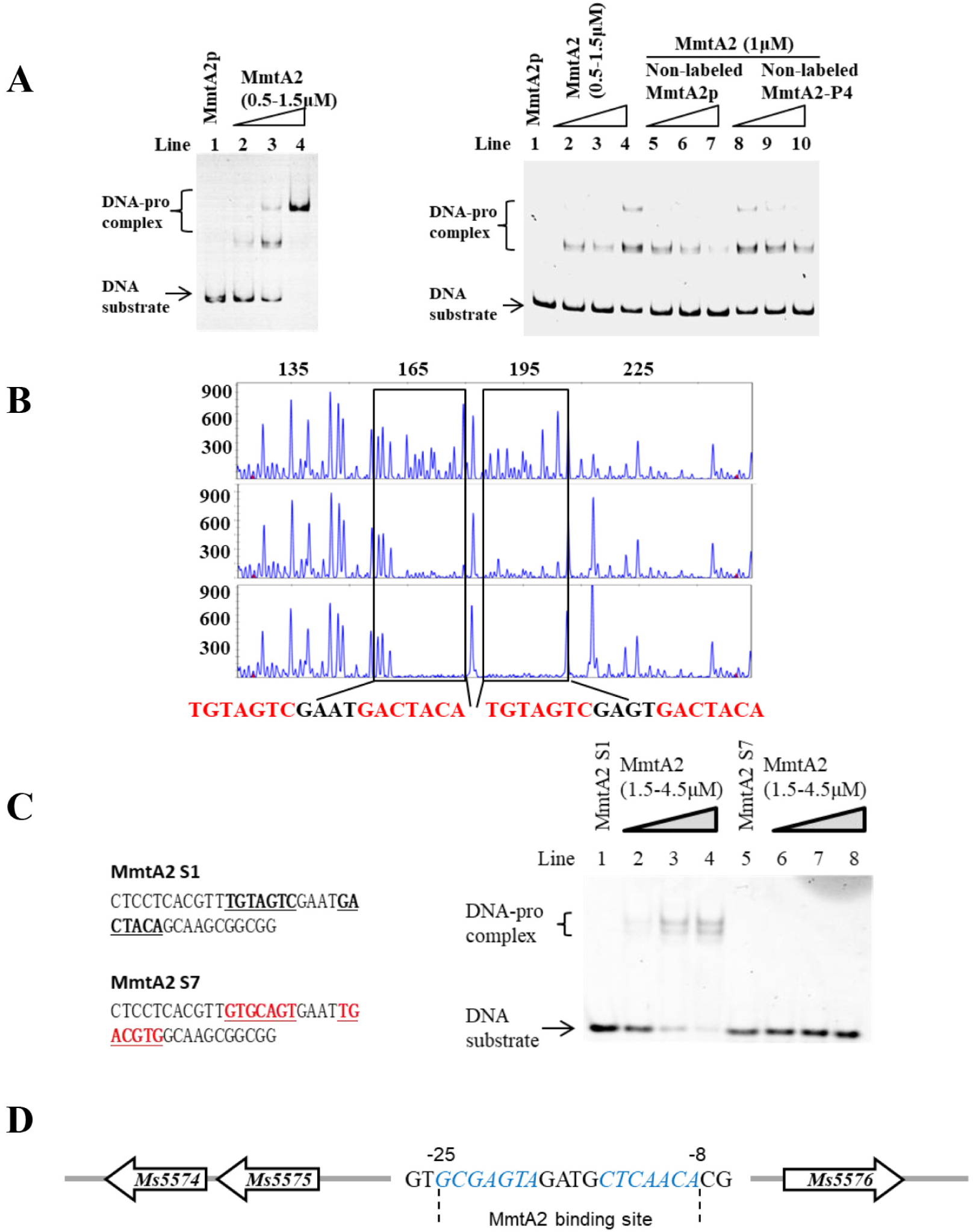
Transcriptional factor MmtA2 binds to a palindromic motif. (A) EMSA analysis of MmtA2 DNA binding activity to MmtA2 promoter DNA. *Left*, MmtA2p (lanes 1–4) and MmtA2-P4 (lanes 5–8) co-incubated with increasing concentrations of purified MmtA2 protein (0, 0.5, 1 and 1.5 μM). *Right*, competitive binding assay for specific DNA-binding activity of MmtA2 on MmtA2p. (B) DNaseI footprinting assays. FITC-labeled DNA fragment was incubated at the absence (*top*) or in the presence of MmtA2 protein (*middle*, 0.5 μM; *bottom*, 1.5 μM). (C) EMSA analysis of MmtA2 binding to DNA substrates with (lanes 1–4) or without (lanes 5–8) identified IR sequences. Either DNA substrate was co-incubated with 1.5–4.5 μM of MmtA2 protein. (D) Hypothetical MmtA2 binding site in the *mmt* promotor.

By using β-galactosidase assays, we found LacZ activity was significantly increased in the Δ*mmtA2* strain when promoted by the MmtA2p promoter (Fig. S3B), suggesting that MmtA2 functions as an auto-repressor through its binding to the motif located inside *mmtA2* promotor. Additionally, a potential MmtA2 binding motif was identified in the *mmt* promotor region, from −22 to −8 bp upstream of the *Ms5576* gene (Fig. 6D).

### MmtA1 positively regulates sugar transportation in *M. smegmatis*

Analysis of the RNA sequencing results (GSE110455) (17) revealed three gene clusters related to sugar metabolism and transportation, including the *mmt* operon (*Ms5571*-*Ms5576*), were up-regulated in both *mmtA1*- and *mmtA2*-overexpressing strains compared to the wild-type strain (Fig. 7A). Further quantitative real-time PCR assays confirmed that both MmtA1 and MmtA2 positively regulated the expression of the gene clusters *Ms3267*-*Ms3269* and *Ms6802*-*Ms6804*. As shown in Fig. 7C, the expression levels of all six genes were significantly increased in the *mmtA1*- and *mmtA2*-overexpressing strains. Comparative sequence analysis suggests that the promotors of these two gene clusters contain MmtA2 binding site (Fig. 7B). The results indicate that MmtA1 regulates several sugar metabolism and transportation pathways in *M. smegmatis* through MmtA2.

**Figure 7.**
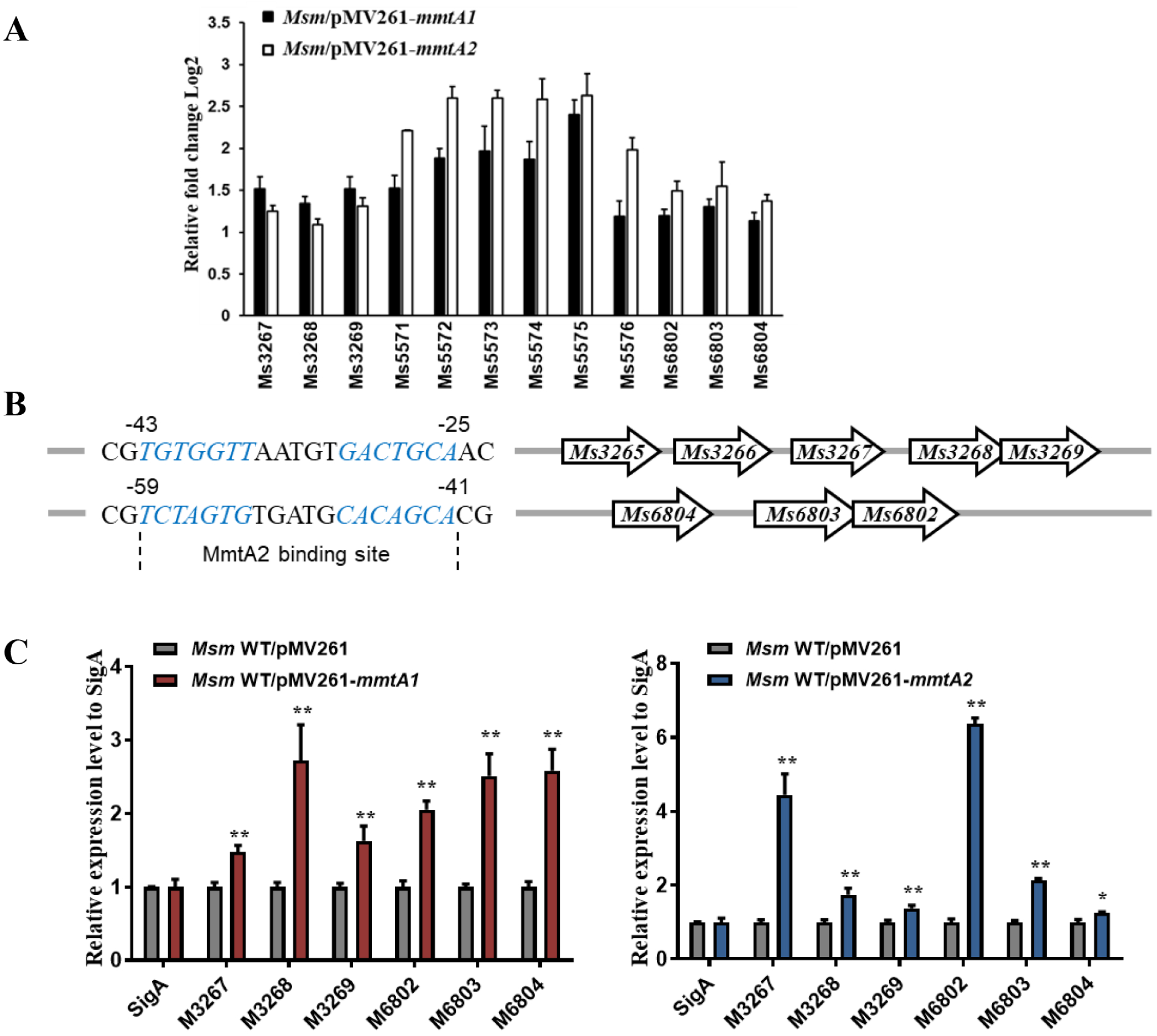
MmtA1 and MmtA2 positively regulate sugar transporter genes. (A) Comparison of differential expression genes in *mmtA1*-overexpressing and *mmtA2*-overexpressing strains compared to the wild-type strain (transcriptomic data, *P*<0.05). (B) Hypothetical MmtA2 binding sites in promotor region of Ms3267-Ms3269 and Ms6802-Ms6804 gene clusters. (C) RT-qPCR assays for target gene mRNA expression levels in *mmtA1*-overexpressing (*left panel*) and *mmtA2*-overexpressing (*right panel*) strains. *, P < 0.05; **, P < 0.01.

### MmtA1 positively regulates lipid transportation in *M. smegmatis*

The RNA sequencing results suggested that 24 of the differentially expressed genes in *mmtA1*-overexpressing strain were enriched in the ABC transporter KEGG pathway (Fig. 8A and Table S2). Compared with the wild-type strain, 13 ABC transporter related genes were up-regulated in the *mmtA1*-overexpressing strain, but not in the *mmtA2*-overexpressing strain. The promoters of these ABC transporter genes contain MmtA1-specific binding sites (Fig. 8B), indicating that MmtA1 directly regulates their expression. Several genes (Ms1141-Ms1142, Ms1144-Ms1148, Ms6540) related to lipid transportation were upregulated in the *mmtA1*-overexpressing strain. Furthermore, GC-MS analysis was used to detect the content of fatty acids in the biofilm of *M. smegmatis* strains. Overexpression of *mmtA1* increased the content of 12-methyltetradecanoic acid in *M. smegmatis* biofilm, while deletion of *mmtA1* decreased its content (Fig. 8C). These results imply that MmtA1 affects biofilm formation by regulating several extracellular substances transportation pathways in *M. smegmatis*, including lipid compounds.

**Figure 8.**
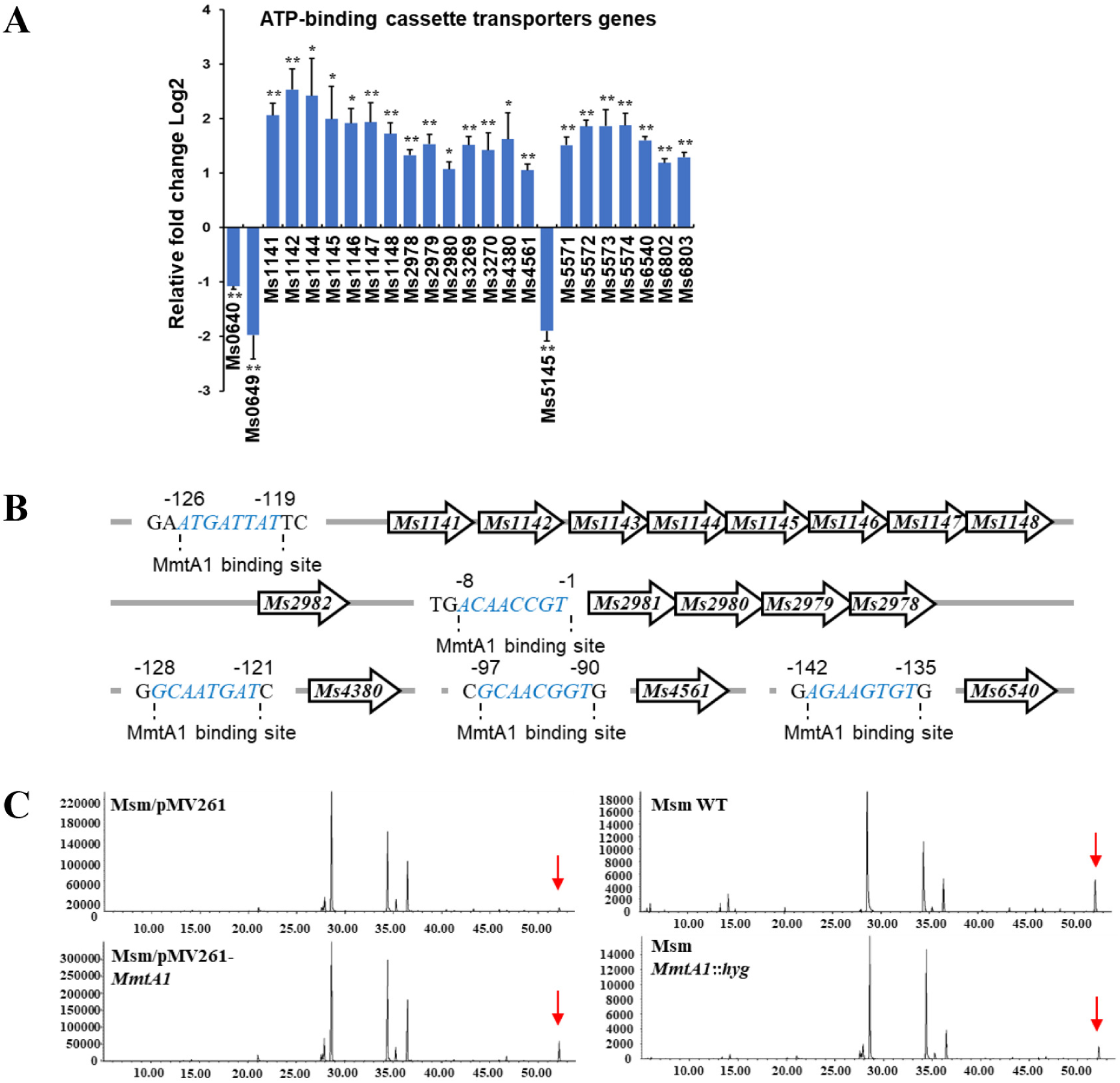
MmtA1 positively regulates lipid transportation in *M. smegmatis*. (A) Relative fold-change of ABC transporter mRNAs in the *mmtA1*-overexpressing strain compared to the wild-type strain. (B) Hypothetical MmtA1 binding site in promotors of target genes. (C) GC-MS analysis of fatty acids in bacterial biofilm. The arrows indicate the compound 12-methyl-tetradecanoic acid, methyl ester.

## Discussion

In this study, we identified a pathway by which antibiotic isoniazid induces multidrug resistance in mycobacteria. Specifically, we found that isoniazid initially up-regulates the expression of the transcriptional factor MmtA1, which then activates the expression of its target genes associated with sugar and lipid transportation. Through this regulation, the target genes promote the biofilm formation and eventually leads to multi-antibiotic resistance in bacteria (Fig. 9). Together with the prior study (17), we proposed a hypothesis that isoniazid may act as a signaling molecule to activate the antibiotic resistance via the transcriptional regulatory pathway governing by MmtA1. Therefore, we show a series of self-protection reactions of mycobacteria at the transcriptional level in response to isoniazid.

**Figure 9.**
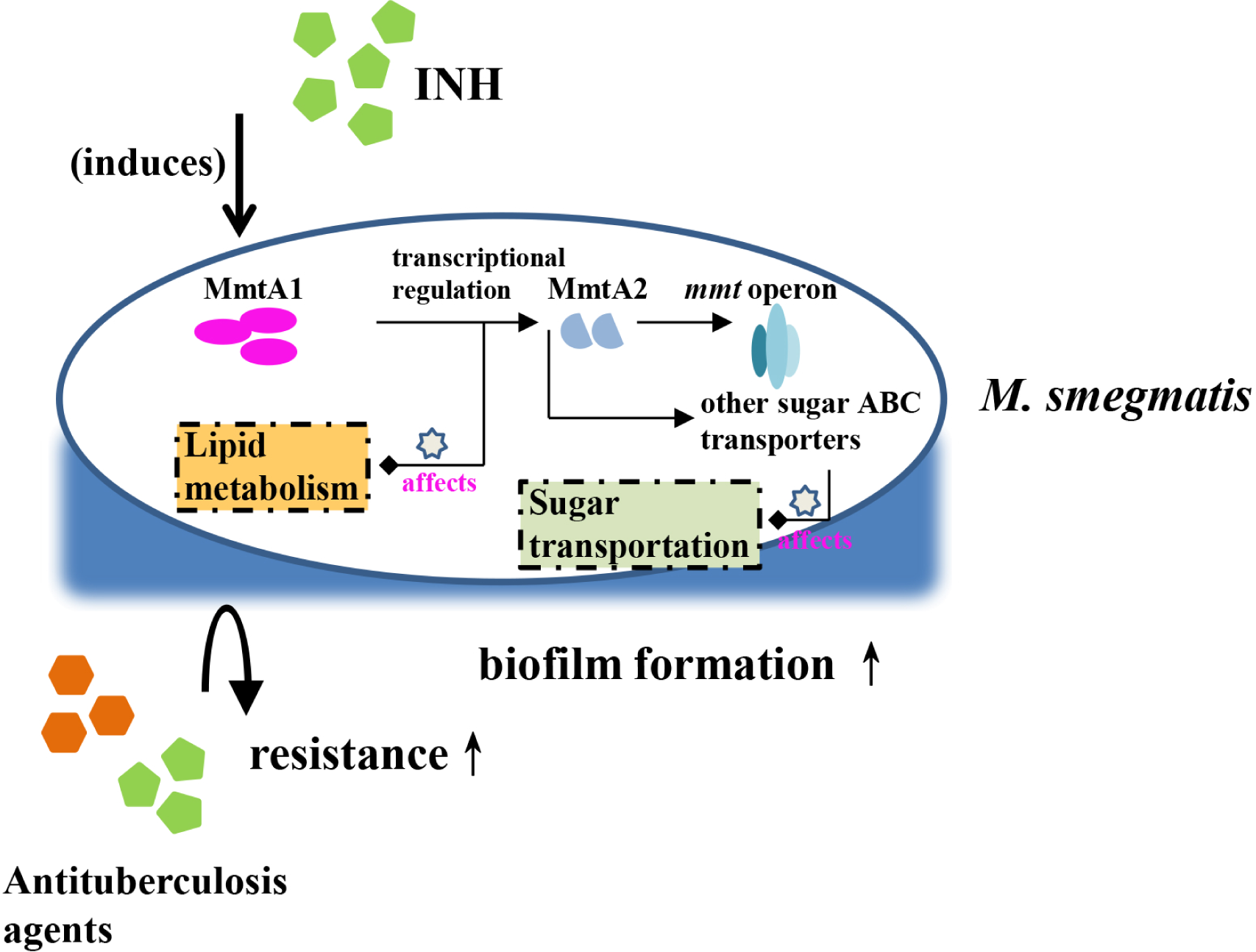
Schematic diagram summarizing the role of transcription factor MmtA1 in regulating drug resistance in *M. smegmatis*. The transcriptional factor MmtA1 which induced by INH, up-regulated target genes in sugar and lipid transportation pathways, leading to multi-drug resistance in *M. smegmatis*.

Mycobacterial biofilms are embedded in a self-produced matrix (EPS) which is mainly composed of exopolysaccharides, lipids, proteins, and extracellular DNA (19). Studies have suggested that extracellular sugar and their metabolism are involved in the biofilm formation of mycobacteria. For example, cellulose has been identified as a structural component of mycobacterial biofilms in both *M. tuberculosis* and *M. smegmatis* (7, 20, 21). As previously demonstrated, MmtA1 regulates the metabolism and transportation of a sugar compound, D-mannitol, and affects biofilm formation in *M. smegmatis* (17). Our results suggest that MmtA1 acts as a sugar ATP-binding cassette (ABC) transporter activator. In bacteria, ABC transporters are required for the import or export of a wide range of substrates, such as sugars, lipids, amino acids, toxins and antimicrobial agents (22, 23). Several ABC transporter genes were previously proved significantly upregulated in *M. tuberculosis* biofilm, indicating ABC transporters are potential target genes of mycobacterial biofilm (24). However, regulation of ABC transporter gene expression were seldomly reported in mycobateria. For example, sequence analysis showed MmtA1 shares a motif (Pfam ID: PF01978) with TrmB family proteins, which can act as global transcriptional regulators to regulate sugar ABC transporters in both archaea and bacteria (25). TrmB is diverse transcriptional factor family harboring with an N-terminal DNA binding domain classified to the wHTH superfamily (IPR036388) and a C-terminal ligand sensing domian (IPR021586) (26). However, MmtA1 only have the wHTH-like DNA binding domain (IPR036388) but not the lignd sensing domain, which implies MmtA1 may share the similar DNA binding proporteis with TrmB TFs and regulate the sugar ABC tranportation genes in *M. smegmatis*, but differ in the ways of environmental stress response.

However, unlike most other bacteria, mycobacterial cells are encapsuled in lipid-rich extracellular matrix when forming biofilms. Previous studies elucidated mycolic acids and fatty acids synthesis process is implicated in biofilm growth in mycobacteria (5, 6, 27). Evidence suggests MmtA1 upregulated a compound of fatty acid, 12-methyltetradecanoic acid, in *M. smegmatis* biofilm. Earlier study showed that clinically isolated strains of methicillin-resistant *Staphylococcus aureus* (MRSA) contained a high proportion of 12-methyltetracanoic acid, suggesting that the increased amount of 12-methyltetradecanoic acid may responsible for the higher resistance of MRSA mutants (28). Thus, the content of 12-methyltetradecanoic acid in biofilm may lead to acquisition of higher antibiotic resistance in *mmtA1*-overexpressing *M. smegmatis*. However, the regulatory function of MmtA1 in 12-methyltetradecanoic acid biosynthesis or transportation progress remains unclear.

The production of biofilm is usually observed under environmental stress conditions and provides protection for resident bacilli from disadvantageous conditions such as antibiotic killing (29). In this study, the biofilm growth of several gene-overexpressing strains may be related to their resistance to the anti-tuberculosis drug isoniazid. Furthermore, strains over-producing transcriptional factor MmtA2 and ABC transporter Ms5571-Ms5573 showed arrested growth at stationary phase when compared with wild-type in normal culture conditions. An implication of this result is that more abundant of biofilm production in these strains block their access to nutrients in environment. It was assumed that while the peripheral biofilm was growing, nutrients supplement for interior cells may be limited due to the increased layers of cells (30). As a result, the growth rate of the bacterial population was also limited. It infers that when exposed in stressed environment, the opposing demands for bacterial growth and maintaining viability of protected interior cells in biofilm was conflict (30). Therefore, the precise control of gene transcription that modulate physiological transitions in progress of biofilm growth during various environmental stress conditions is of great significance.

### Conclusions

Our study illustrated a self-protection mechanism through regulatory pathways in mycobacteria when exposed in antibiotic environment, implying that in addition to act as antibiotic, isoniazid may be involved in the regulation of biofilm formation as a molecular messenger, leading to multiple-drug resistance of *M. smegmatis*. This finding provides a novel insight into drug-induced resistance mechanism of mycobacteria under antibiotic drug pressure.

## Materials and Methods

### Bacterial strains, plasmids, enzymes, and agents

E. coli BL21 (λ DE3) cells and pET28a plasmids from Novagen (Darmstadt, Germany) were used for recombinant proteins expression. *M. smegmatis mc2 155* strains was grown in 7H9 Middlebrook medium (Becton Dickinson, USA) supplemented with 0.2% glycerol and 0.05% Tween 80. The enzymes and all antibiotics were from TaKaRa Biotech (Shiga, Japan). The construction methods of the gene-overexpressing and knockout *M. smegmatis* strains were described in our previous work (17).

### Cloning, expression, and purification of recombinant proteins

The genes *mmtA1* and *mmtA2* were amplified from *M. smegmatis* genome by using specific primers (Table S1). The PCR products were purified and cloned into the pET28a vector. The fusion-expression plasmids were then transformed into *E. coli* BL21 (DE3) cells to express target protein. 0.5 mM IPTG (isopropyl β-D-1-thiogalactopyranoside) was used to induce the expression of the *mmtA1* and *mmtA2* genes. The His-tagged fusion proteins were affinity purified (Ni-NTA agarose resin; Qiagen, Germany) and quantitated as described previously (31).

### Minimum inhibitory concentration (MIC) determination

The MIC of isoniazid on *M. smegmatis* strains was evaluated by the agar diffusion procedure (32). *M. smegmatis* strains were pre-inoculated onto 7H10 agar plates (150 mm), then filter disks (10 mm) with 0 to 7 ng isoniazid were applied to the inoculated agar. The plates were incubated at 37 ℃ for 48 hours before measuring the diameters of discernible inhibition zones. The minimum inhibitory concentrations of isoniazid to strains were then calculated.

### Determination of bacterial viability on the effect of antibiotics

Growth patterns of the mycobacterial strains were examined according to the procedures described previously with some modifications (33). *M. smegmatis* strains were first grown in Middlebrook 7H9 medium at 37 ℃ while shaking at 160 rpm. Cells were cultured into a stationary phase and diluted in 100 ml of fresh 7H9 broth (OD_600_ ≈ 0.15). *M. smegmatis* cultures were treated with 15 μg/ml isoniazid, 6 μg/ml rifampicin or 15 μg/ml isoniazid compared with untreated controls to determine antibiotic effect on bacterial viability. Aliquots were taken at the indicated times, *M. smegmatis* colonies were measured and CFU was calculated. The experiments were carried out in triplicate.

### RNA extraction and quantitative real-time PCR

The *M. smegmatis* cells were first culrured in 7H9 fluid medium at 37°C. When bacteria reach the stationary phase, various concentration of isoniazid was added into the medium. After co-incubated with antibiotic for 3 h, cells were harvested. Then total RNA was extracted with Qiagen RNeasy Mini kit according to the manufacturer’s recommendation (34).

Primers used for the real-time PCR were listed in Table S1. Real-time quantitative PCR was performed as previously described (33). Expression levels of candidate genes were normalized to the levels of *sigA* gene transcripts. As proved previously [31], the gene *ms0535* not induced by isoniazid was used as a negative control. Gene expression variation presented by using a modification of the 2^−ΔΔCt^ method. Differential expression was reported as mRNA fold change. Student *t*-test was used and a *P* value ≤0.05 was considered to be statistically significant.

### Quantitative analysis of *M. smegmatis* biofilm

The crystal violet (CV) assay for quantification of bacterial biofilm formation was performed as described previously (17). Prior to inoculation, strains were cultured with 7H9 broth media to the stationary phase. The strains were then collected and transferred to M63 minimal media supplemented with 1 mM MgSO_4_ and 22 mM glucose, the cell suspension was inoculated into a 96-well PVC microtiter plate with 100 μL per well. The inoculated plate was incubated at room temperature for 24 h with slightly shaking at 80 rpm. After associated with CV staining, the biofilm was extracted with ethanol/acetone (v/v = 80:20) and measuring the OD at 570 nm using a TecanNanoQuant microplate reader (Männedorf, Swiss).

### Electrophoretic mobility shift assay

Electrophoretic mobility shift assays (EMSA) were performed to determine the binding capacity of MmtA1 and MmtA2 to their operator regions. The target DNA fragments MmtA1p, MmtA2p and negative controls were amplified from *M. smegmatis* genomic DNA with specific primers (Table S1). Experiments were conducted as previously described (33). To subsequently test for the binding specificity of transcriptional factors, promoter substrates were amplified by primers carry a 5’-label of FITC (33). The protein combined 5’-FITC DNAs were incubated with increasing concentrations of unlabeled promoter DNA or non-specific competitors. Signals were acquired by using a Typhoon Scanner (GE Healthcare).

### DNase I Footprinting

For DNaseI footprinting assays, the indicated amount of candidate proteins was incubated in a 100 μL reaction containing approximately 2.5 nM of fluorescently labeled DNA in reaction mixtures at 37 ℃ for 30 min (35). 5 U of DNase I (TAKARA) was added and incubated for 1 min. The digestion was terminated with the addition of equivoluminal phenol/chloroform (phenol:chloroform:isopropanol = 25:24:1) extraction. After ethanol/ sodium acetate precipitation and 70% ethanol washing, the DNA was resuspended in 10 μL of TE buffer. The 500-bp fragments were sequenced using the Thermo Sequenase Dye Primer Manual Cycle Sequencing Kit (USB, Inc., Cleveland, OH, USA) according to the manufacturer’s instructions. Capillary electropherograms were analyzed and aligned by the GENEMAPPER software (version 4.0, Applied Biosystems) (33).

### Analysis of β-galactosidase activity

β-galactosidase activity assays were performed by constructing promotor-lacZ fusion plasmids derived from pMV261 (36). The promotor DNA fragments were obtained by amplification with specific primers (Table S1). The recombinant plasmids were transformed into wild-type or knockout *M. smegmatis* to generate reporter strains. The strains were pre-cultured in a 7H9-Tw-glycerol-Kan liquid medium and grown to the stationary stage. After harvested, β-galactosidase activity of the different strains was measured and presented as Miller units.

### Determination of fatty acids in M. smegmatis biofilm

*M. smegmatis* cells were harvested after 24 hours of growth in Middlebrook 7H9 medium. Bacterial biofilm was lyophilized prior to lipid extraction and total lipids were extracted with 15% NaOH/MeOH solution at 55 ℃ for 3 h. After the reaction, the solution was neutralized with an appropriate amount of HCl, and esterified fatty acids were extracted by hexane acetate (37). The GC-MS assay was applied to analyze the fatty acid composition by using Agilent 7890A gas chromatograph coupling with an Agilent 5975C mass spectrometer. The chromatographic column used for analysis was HP-5MS column (5% Phenyl Methyl Silox 30 m×250 μm×0.25 μm).

## Funding

This research was funded by the China Postdoctoral Science Foundation, grant number 2019M662730.

### Data Availability Statement

Data are contained in this manuscript or Supplementary Materials. The RNA sequencing data have been submitted to NCBI.

## Acknowledgments

We especially thank professor Zheng-Guo He for the support of experimental material and advice on experimental design.

## Conflicts of Interest

The authors declare no conflict of interest.

